# Priming winter wheat seeds with the bacterial quorum sensing signal N-hexanoyl-L-homoserine lactone (C6-HSL) shows potential to improve plant growth and seed yield

**DOI:** 10.1101/490391

**Authors:** Olena V. Moshynets, Lidia M. Babenko, Sergiy P. Rogalsky, Olga S. Iungin, Jessica Foster, Iryna V. Kosakivska, Geert Potters, Andrew J. Spiers

## Abstract

Several model plants are known to respond to bacterial quorum sensing molecules with altered root growth and gene expression patterns and induced resistance to plant pathogens. These compounds may represent novel elicitors that could be applied as seed primers to enhance cereal crop resistance to pathogens and abiotic stress and to improve yields. We investigated whether the acyl-homoserine lactone N-hexanoyl-L-homoserine lactone (C6-HSL) impacted winter wheat (*Triticum aestivum* L.) seed germination, plant development and productivity, using two Ukrainian varieties, Volodarka and Yatran 60, in both *in vitro* experiments and field trials. *In vitro* germination experiments indicated that C6-HSL seed priming had a small but significant positive impact on germination levels (1.2x increase, p < 0.0001), coleoptile and radicle development (1.4x increase, p < 0.0001). Field trials over two growing seasons (2015-16 and 2016-17) also demonstrated significant improvements in biomass at the tillering stage (1.4x increase, p < 0.0001), and crop structure and productivity at maturity including grain yield (1.4 – 1.5x increase, p < 0.0007) and quality (1.3x increase in good grain, p < 0.0001). In some cases variety effects were observed (p ≤ 0.05) suggesting that the effect of C6-HSL seed priming might depend on plant genetics, and some benefits of priming were also evident in F1 plants grown from seeds collected the previous season (p ≤ 0.05). These field-scale findings suggest that bacterial acyl-homoserine lactones such as C6-HSL could be used to improve cereal crop growth and yield and reduce reliance on fungicides and fertilisers to combat pathogens and stress.

## Introduction

Wheat is one of the most important food staples and export commodities in Ukraine with ∼95% of the harvest coming from winter wheat crops [1]. Winter wheat represents ∼95% of the annual crop with the remaining 5% planted as spring wheat [2]. It is typically sown in autumn (September-October) and harvested in mid-summer (July) of the following year, however, weather conditions severely affect the quality of the crop, and frequent changes during the growing season makes it more susceptible to bacterial and fungal pathogens than spring wheat which is grown from April to August. While there are a number of intensive agricultural technologies to improve crop yield and control pathogens, the economics of wheat production is dominated by the need to keep inputs beyond pre-plant bactericide and fungicide treatment and fertilization low to be economical. Therefore, there is a need to augment existing production practices with economical, yet effective seed treatments to increase crop yield and resistance to pathogens.

The large-scale use of agricultural bactericides and fungicides, their persistence and impact on the wider environment is of increasing concern, with over 284,411 tonnes used worldwide in 2015 [3]. Their use can be reduced by using plant growth-promoting bacterial inoculants as well as bacterially-derived plant growth regulators to treat seeds and seedlings [4-6]. Such ‘green’ technology is becoming more popular, and in some cases, natural phyto-stimulators provide a more lasting impact on crop stress tolerance and increase productivity without undesirable environmental effects [7].

One of the most effective technologies to increase crop resistance to biotic and abiotic stressors is seed priming which increases germination vigour and activates plant defence mechanisms early in plant development through induced resistance [8,9]. Plant growth can be further supported by bacterial inocula which can facilitate mineral up-take and reduce pathogens through biological competition as well as stimulate the plant defence system [10,11]. Plants are also known to respond to bacterial quorum sensing signal molecules used by bacteria to regulate gene expression and coordinate social activities [8-9, 12-14]. For example, the legume *Medicago truncatula* responds to nanomolar concentrations of acyl-homoserine lactones (AHLs) produced by its symbiont *Sinorhizobium meliloti* and the pathogen *Pseudomonas aeruginosa* with specific and extensive changes in root gene expression and protein accumulation [15]. In tomato (*Solanum lycopersicum*) the plant growth-promoting bacterium *Serratia marcescens* but not AHL-deficient mutants increase systemic resistance against the fungal leaf pathogen *Alternaria alternata* [16], and treatment of roots with synthetic AHLs enhanced the expression of defence-related genes in the leaves of tomato [17].

We are interested in determining whether bacterial AHLs used as a seed primer can have a significant impact on winter wheat crop yields under realistic agricultural conditions. Here we compare plant development, productivity and yield structure for two Ukrainian winter wheat varieties following seed priming with the AHL N-hexanoyl-L-homoserine lactone (C6-HSL) and demonstrate improved crop yields over two growing seasons in 2015 – 17 at the Feofaniya research site near Kiev.

## Materials and methods

### C6-HSL synthesis

L-homoserine lactone hydrochloride was synthesized using a modified procedure [18] (**Fig 1**). 30 g of L-methionine, 120 mL water, 120 mL isopropanol and 50 mL glacial acetic acid were added to a 1 L round-bottom flask equipped with a magnetic stirrer. 20.8 g chloroacetic acid added and the suspension stirred for 24 h at 70°C. The resulting transparent solution was boiled with reflux for 3 h and the solvents distilled off at a reduced pressure. To convert the homoserine into its lactone, the solid residue was heated for 6 h at 100 – 110°C and reduced pressure of 15-20 mbar. The product was dissolved in 50 mL water before 250 mL isopropanol was added and precipitated overnight at 4°C. The crystalline solid was filtered out, washed with 100 mL isopropanol and vacuum–dried for 12 h at 40°C and 1 mbar. In this procedure the yield of L-homoserine lactone hydrochloride was 65%. A proton nuclear magnetic resonance (^1^H NMR, 300 MHz, DMSO-D6) spectrum of the end product with the following characteristics was obtained: ^1^HNMR (DMSO-D6): δ 2.37 (^1^H, m, 4α-H), 2.56 (^1^H, m, 4β-H), 4.31 (^2^H, m, 5α-H, 3-H), 4.34 (1H, td, 5β-H), 9.05 (3H, br s, NH).

**Fig 1.**
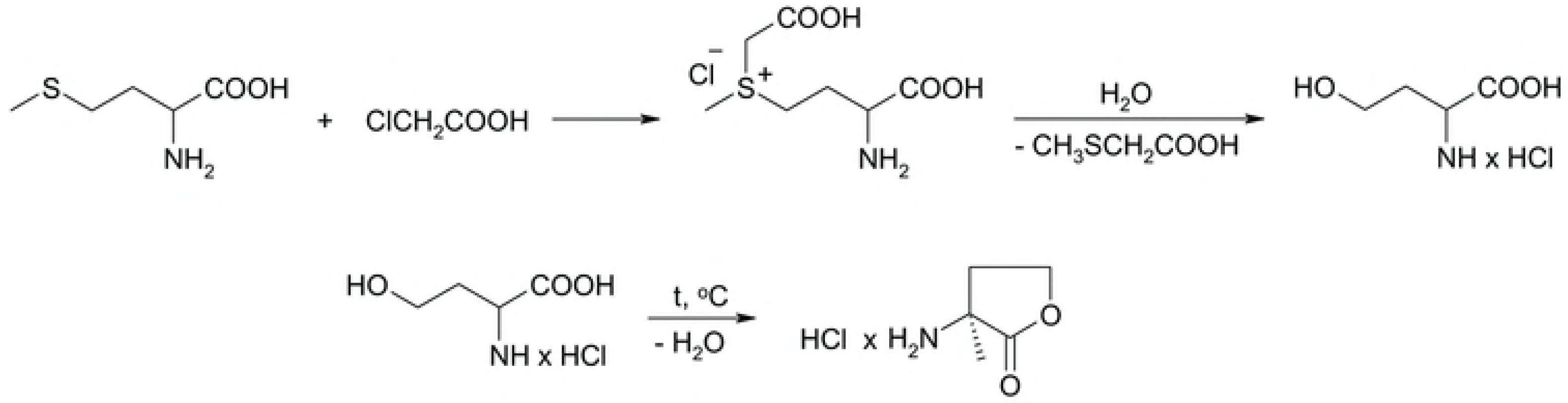
Synthetic route to obtain L-homoserine lactone hydrochloride.

The second-step synthesis of N-hexanoyl-L-homoserine lactone (C6-HSL) followed an earlier method [19] (**Fig 2**). 8.7 g trimethylamine was added to a stirred suspension of 5 g L-homoserine lactone hydrochloride in 70 mL methylene chloride at 0°C. The mixture was stirred for 30 min before 4.9 g hexanoyl chloride was added dropwise within 10 min. The resultant reaction continued for 1 h at 0 – 5°C and then at room temperature for a further 4 h. The mixture was washed twice with 50 mL saturated sodium hydrogen carbonate, twice with 50 mL 1 M potassium hydrogen sulfate, and then with 70 mL saturated sodium chloride. The solution of C6-HSL in methylene chloride was dried over sodium sulfate and the solvent was removed at reduced pressure. The white solid residue was further purified by double-recrystallization in ethyl acetate (1 g of product per 15 mL). In this procedure the yield of C6-HSL was 60%. A proton nuclear magnetic resonance (^1^H NMR, 300 MHz, DMSO-D6) spectrum of the end product with the following characteristics was obtained: ^1^HNMR (DMSO-D6): δ 0.89 (3H, t, CH_3_), 1.31 (4H, m, CH_3_(CH_2_)_2_), 1.64 (2H, m, CH_2_CH_2_CO), 2.17 (1H, m, 4 α-H), 2.25 (2H, t, CH_2_CO), 2.79 (1H, m, 4 β-H), 4.29 (1H, m, 5α-H), 4.46 (1H, td, 5 β-H), 4.64 (1H, m, 3-H), 6.5 (1H, d, NH).

**Fig 2.**
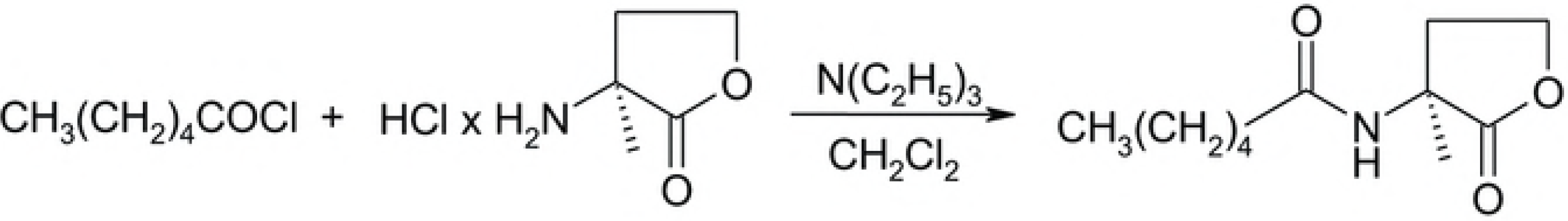
Synthetic route to obtain N-hexanoyl-L-homoserine lactone.

### Winter wheat varieties

Two varieties of winter wheat (*Triticum aestivum* L.), Volodarka and Yatran 60, cultivated extensively in the Lisostep and Polissye regions of Ukraine and produced by the Institute of Plant Physiology and Genetics (National Academy of Science of Ukraine) and the V.M. Remeslo Mironovka Institute of Wheat (Ukrainian Academy of Agrarian Sciences) were used in this work. Volodarka is a short-stalked variety with high yields and good resistance to frost and drought while Yatran 60 is a short-stalked, medium-early variety with good resistance to heat and drought.

### Seed preparation

Seeds were primed in a 100 ng/mL C6-HSL solution (at 0.43 kg/L) for 3 h before being dried for 72 h at 24 – 25°C to a water content of 12 – 14% (w/w). Non-primed control seeds were treated with water before drying. Seeds were then coated using AREAL BS-05 (Legion Company Ltd, Ukraine) pelleting mixture before sowing. Seeds were collected from plants grown from primed seeds in 2015-16 (F1 seeds) and used in further tests.

### Seed germination assays

The effects of seed priming on germination were investigated according to the International Seed Testing Association (ISTA) standards. Fifteen samples of 20 seeds of each variety and treatment were germinated in the dark at 24°C for 24 h in Petri dishes containing Knop’s nutrient solution [20] to determine germination levels. In parallel, three samples of 20 seeds of each variety and treatment were germinated on wet filter paper and coleoptile and radicle lengths determined after 2 days.

### Crop trials

Winter wheat crops were planted and harvested over two consecutive growing seasons (September 2015 – July 2016, and August 2016 – July 2017) at the Feofaniya research site (M.G. Kholodny Institute of Botany, National Academy of Science Ukraine) near Kiev. The crop fields are composed of grey forest soil with ash and a very loamy chernozem (pH KCl 5.7, total humus content 4.1%, 138 mg phosphorus/kg, 90 mg potassium/kg). Seeds were planted at a density of 600 seeds/m^2^ at 3.5 – 4 cm depth. Nitrogen, phosphorus and potassium fertilizer was added during the growing season. Weather conditions (**Fig 3**) were recorded by the Meteorological Centre, Central Geophysical Observatory of B.I. Sreznevsky at Kiev, approximately 6 km from Feofaniya.

**Fig 3.**
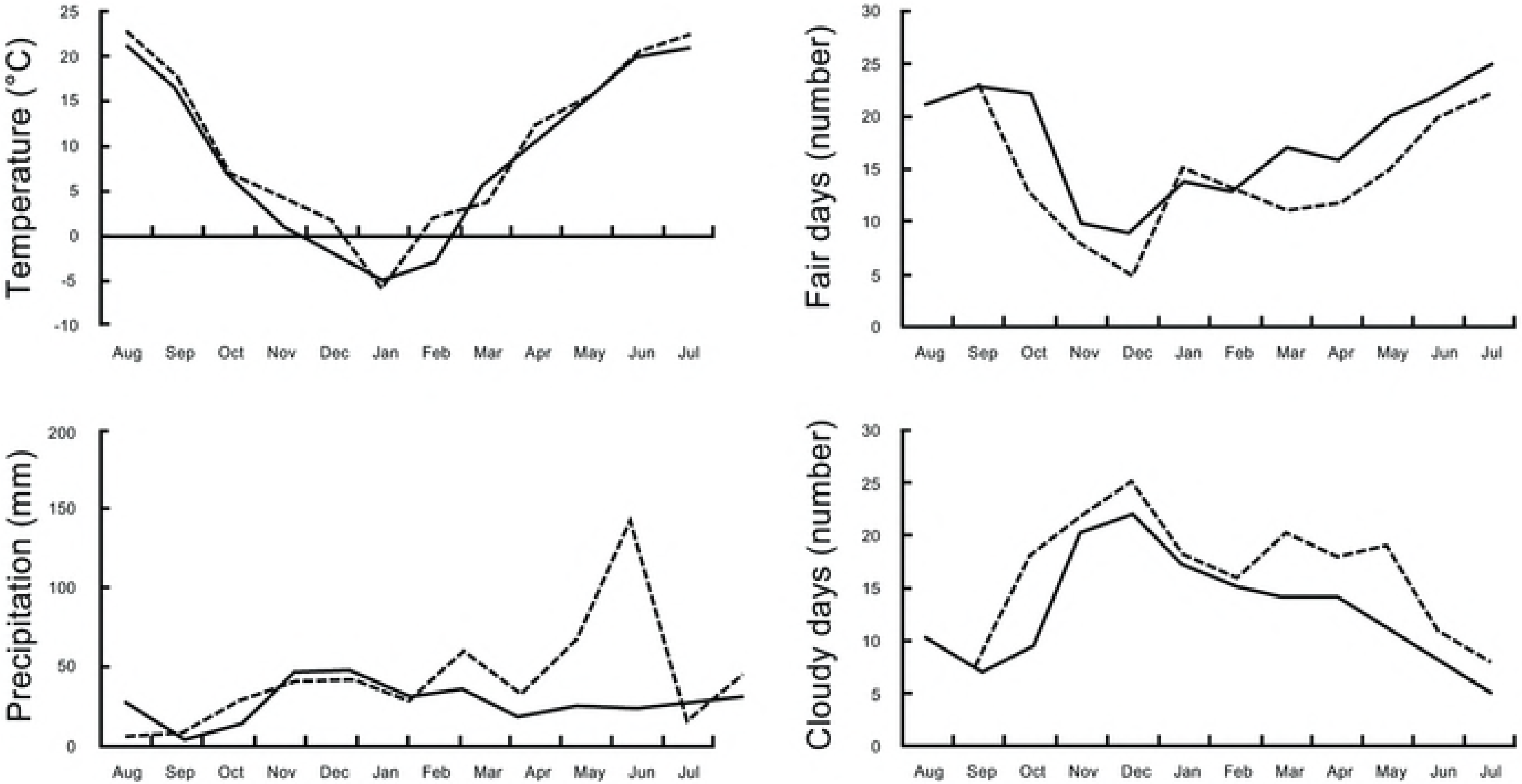
Meteorological conditions during crop trials. Winter wheat was planted and harvested over two consecutive growing seasons, from September 2015 – July 2016 (dashed lines) and August 2016 – July 2017 (solid lines) at the Feofaniya research site and weather conditions recorded at the nearby Meteorological Centre of the Central Geophysical Observatory of B.I. Sreznevsky.

Morphological parameters describing plant and seed yield were measured for 100 plants of each variety in 2015 – 2016 and 50 plants per variety in 2016 – 2017. Seed quality was assessed visually as either ‘normal’ or ‘poor’ and the proportion of normal seeds determined for three samples of 100 seeds. Five samples of 10 seeds of each type were weighed to obtain mean seed weights. Plants were sampled to assess growth, plant morphology, and seed yield.

### Chlorophyll and carotenoid levels

Winter wheat crops were planted as for the crop trials described above (August 2016 – July 2017) at the Hlevakha research site (Institute of Plant Physiology and Genetics, National Academy of Science of Ukraine) approximately 15 km away from Feofaniya and having a similar soil type. Chlorophyll (a and b) and carotenoid levels were measured after acetone extraction [21]. Five extracts were prepared from material harvested from five plants at each of the booting, flowering, milk ripeness, and milk-wax ripeness stages.

### Microbiological characterization

The numbers of rhizosphere-associated chemo-organotrophic, nitrogen-fixing, denitrifying, and amylolytic bacteria adhering to wheat roots were determined using viable colony counts. Roots from 125 2 month-old plants from 2015-2016 were washed in sterile 0.9% (w/v) saline and aliquots of serial dilutions spread onto Nutrient Agar (HiMedia, India) which were incubated at 24°C for 5 days to determine chemo-organotrophic bacterial numbers. Ashby’s medium supplemented with 2% (w/v) sucrose [22-23] was used to determine nitrogen-fixing bacterial numbers with colonies re-isolated three times in liquid medium to distinguish nitrogen-fixers from oligonitrophilic bacteria [22]. Nitrifying bacteria were isolated in liquid Watson and Mendel’s media and incubation for 20 days [22]. Denitrifying bacteria were isolated using Gil’tai’s medium (pH 6.8 – 7.2) in an inert argon atmosphere using a Hangeit-like technique [24] at 24°C for 5 days, with viable numbers determined using the most probable number method [25] with 3-tube variation [26]. Denitrifying bacteria were identified based on medium alkalinisation, increasing optical density, gas production, and the absence of nitrate. Amylolytic bacterial numbers were determined using 2% (w/v) starch plates incubated at 28°C for 3 days washed with 0.1% (w/v) iodine to visualize cleared halo around the colonies [27].

### Statistical analyses

Data were analysed by modelling using a least squares approach with effects (year, plot, wheat variety, seed treatment, and experimental replicate) tested by LSMeans Differences Student’s t and Tukey HSD tests (α = 0.05) (JMP v12, SAS Institute Inc., USA). Chi-Square tests of independence (www.socscistatistics.com) were used to assess the effect of seed treatment on rhizosphere bacteria. Means and standard errors (SE) are shown where appropriate.

## Results

### Germination

*In vitro* germination assays were conducted to determine whether seed priming effected germination and seedling development (**Fig 4**). Priming had a significant positive effect on germination with levels 1.2x greater than the untreated controls (p < 0.0001), and on two-day old seedling development with coleoptile (the protective sheath covering the emerging shoot) and radicle (embryonic root) lengths 1.4x greater than the untreated controls (p < 0.0001). However, no wheat variety effects were observed in these assays (p = 0.32, 0.99 and 0.48, respectively). These findings suggest that seed priming with C6-HSL may have a positive impact on the early development of wheat seeds.

**Fig 4.**
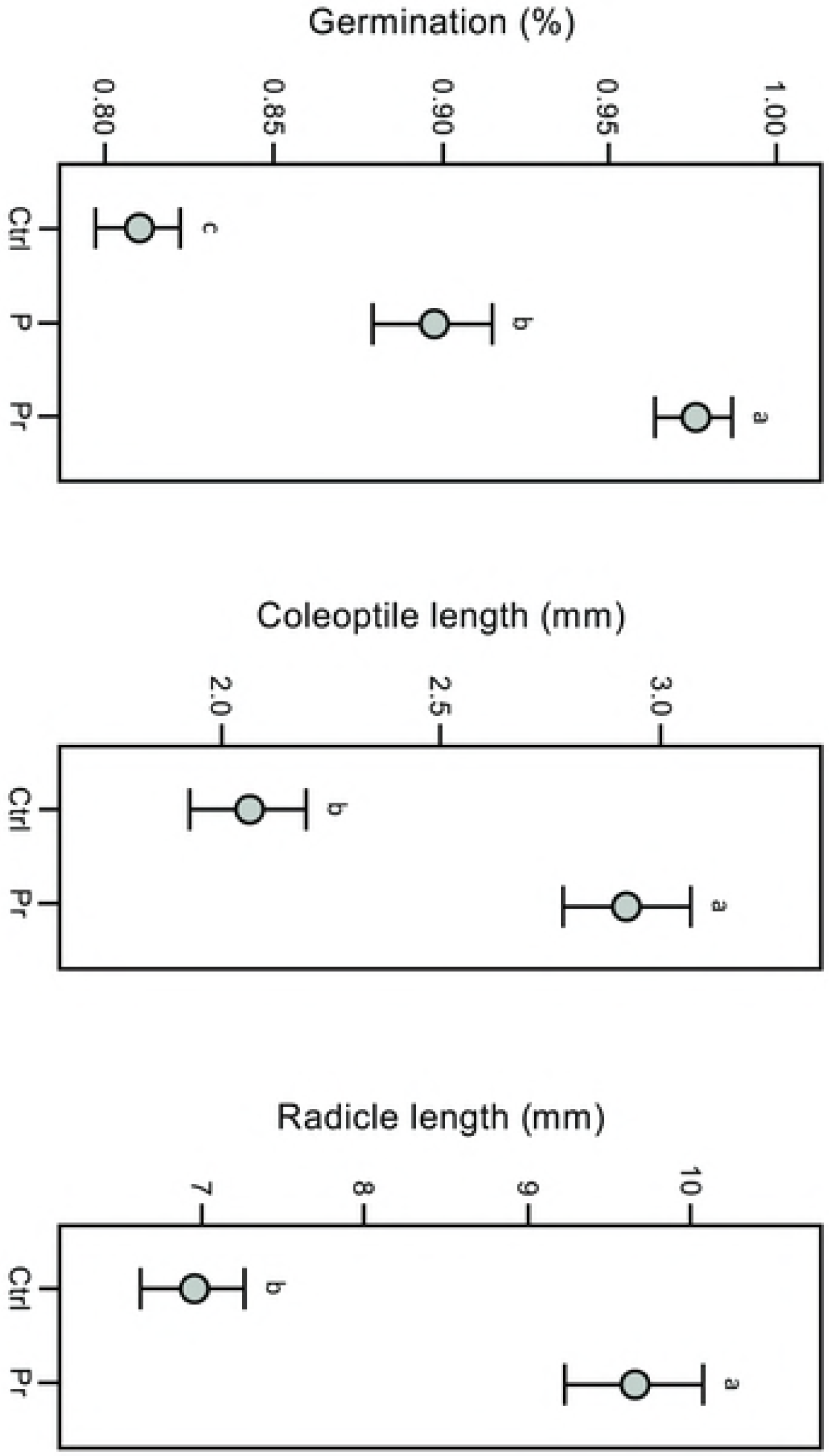
C6-HSL seed priming affects germination and early development. Seed treatments have small but significant effects on *in vitro* germination rates and on 2 day-old winter wheat seedling coleoptile (the protective sheath covering the emerging shoot) and radicle (embryonic root) lengths. Plants were grown from pelleted and primed (Pr), pelleted (P), or untreated (Ctrl) seeds. As no significant wheat variety effect was observed (p = 0.32, 0.99 and 0.48, respectively) data are pooled and means and standard errors (SE) shown. Means not connected by the same letter are significantly different (α = 0.05).

### Plant development

2 month-old plants at the tillering stage were sampled from our research field site in 2015-16 to determine whether seed priming effected plant development (**Fig 5**). Priming had a significant positive effect on plant growth with dried aboveground biomass 1.4x greater than the untreated controls (p < 0.0001), though like the *in vitro* assays, no wheat variety effects were observed (p = 0.39). This suggests that the seed priming advantage seen in better germination rates and faster embryo development also occurs in the field and continues through tillering development.

**Fig 5.**
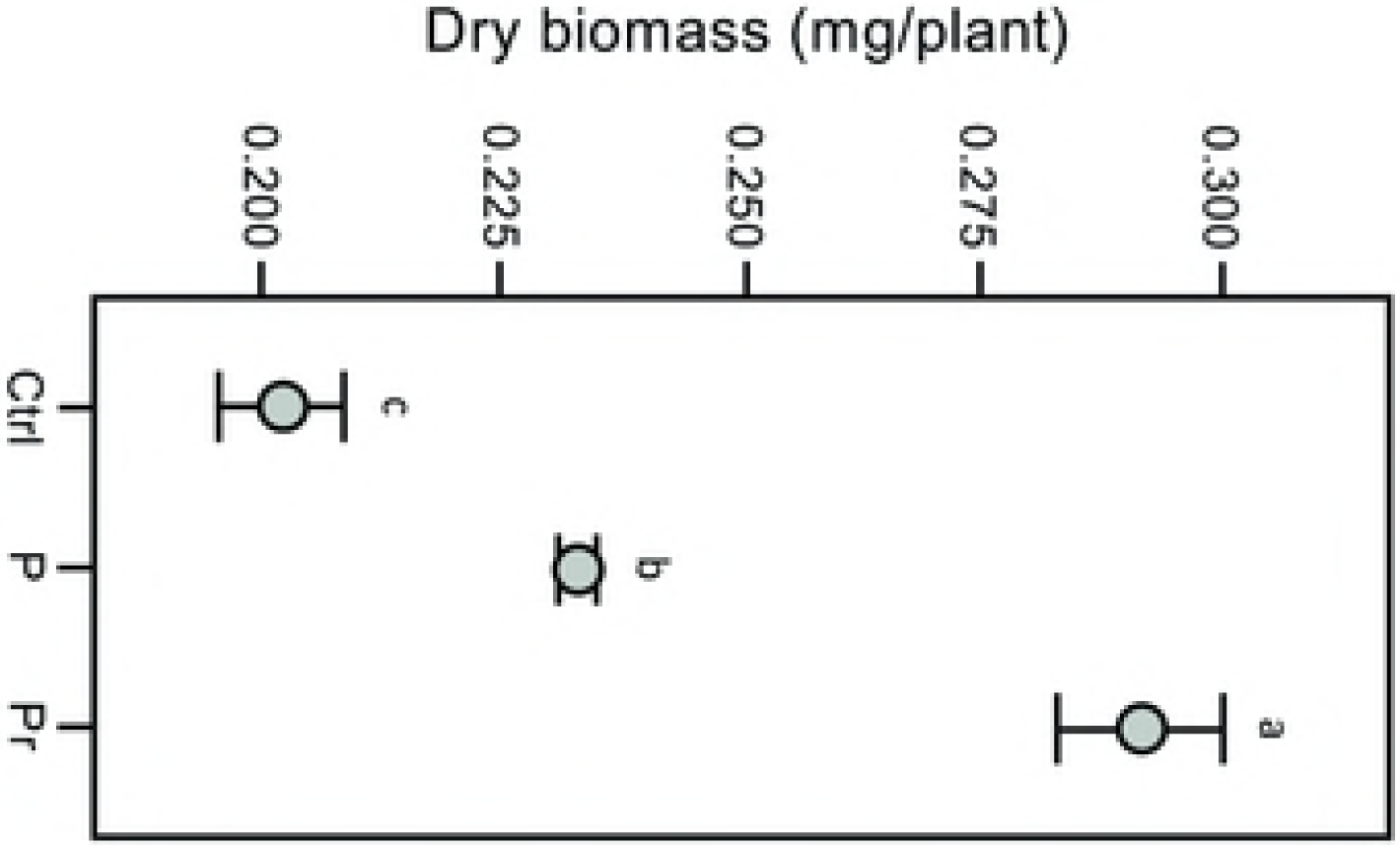
C6-HSL seed priming affects plant development at the tillering stage. Seed treatments have small but significant effects on total dry biomass at 2 months in winter wheat crop trials (2015-16). Plants were grown from pelleted and primed (Pr), pelleted (P) or untreated (Ctrl) seeds. As no significant wheat variety effect was observed (p = 0.3924) data are pooled and means and standard errors (SE) shown. Means not connected by the same letter are significantly different (α = 0.05).

### Crop structure and productivity

Mature plants with grain were harvested in mid-summer of 2015-16 and 2016-17 to determine whether seed priming effected crop structure and productivity. In our analysis of these data, a significant year effect was observed (p < 0.05) which may be explained by the colder conditions and lower seasonal rainfall in the second growing season (**Fig 3**). Priming had a significant effect on over-all plant height, tiller numbers, ear length, spikelets per ear, grains per ear, and the grain mass per plant in 2015-16 and 2016-17 (p < 0.0001), and a significant wheat variety effect was observed (p < 0.0001) with Yatran 60 generally out-performing the Volodarka variety (data for 2015-16 and 2016-17 is provided in **Table 1**). Priming also had a significant positive effect on crop yield (kg/m^2^) with grain mass 1.4x and 1.5x greater than the untreated controls (p < 0.0007) in 2015-16 and 2016-17, respectively. The impact of seed priming with C6-HSL on crop yield is also evident by PCA of data by year and variety (**Fig 6**). In a separate trail at the nearby Hlevakha research site in 2016-17, priming was also found to have a significant effect on over-all chlorophyll and carotenoid levels, as did plant development stage (from booting, flowering, milk ripeness to milk-wax ripeness) (p < 0.0001), and for carotenoids a significant wheat variety effect was also observed (p = 0.006) (**Table 2**). These findings suggest that the positive effect seen with seed priming with C6-HSL continues through wheat development to plant maturity and grain production.

**Table 1.**
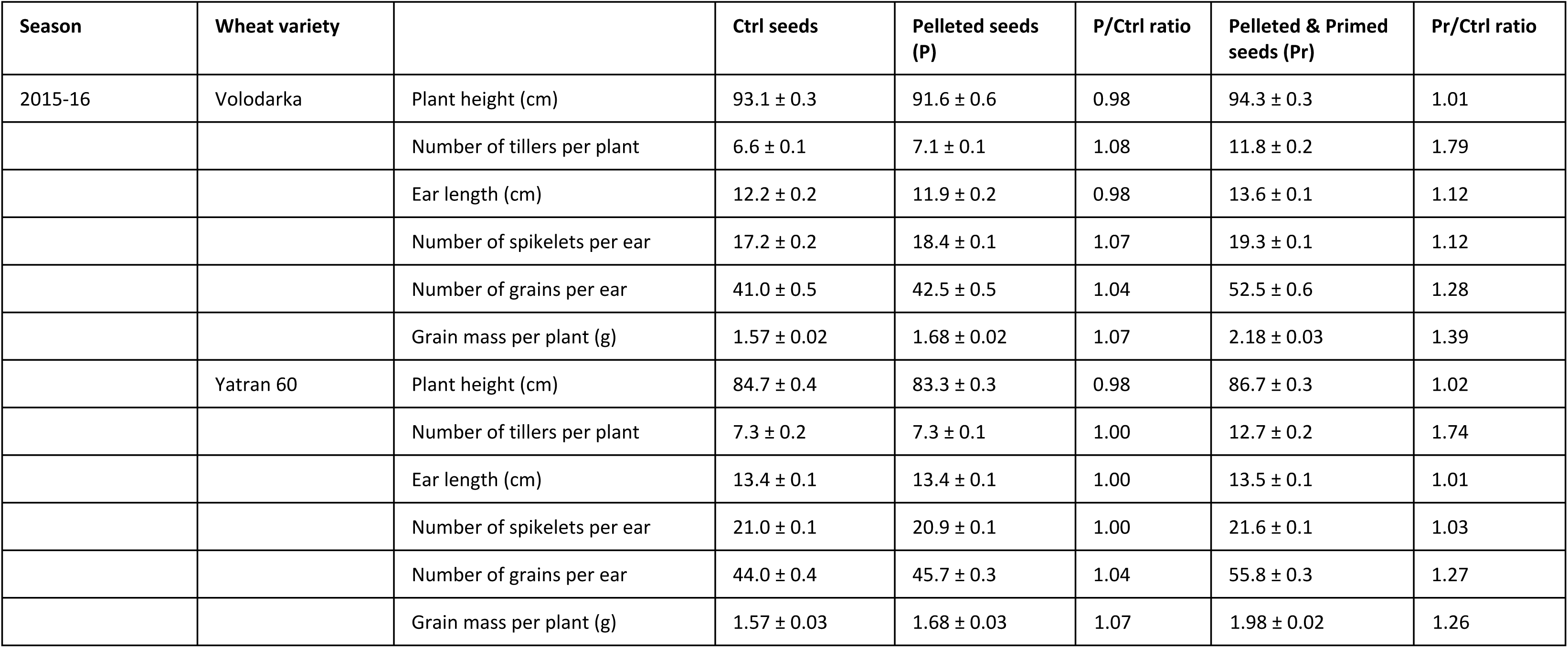
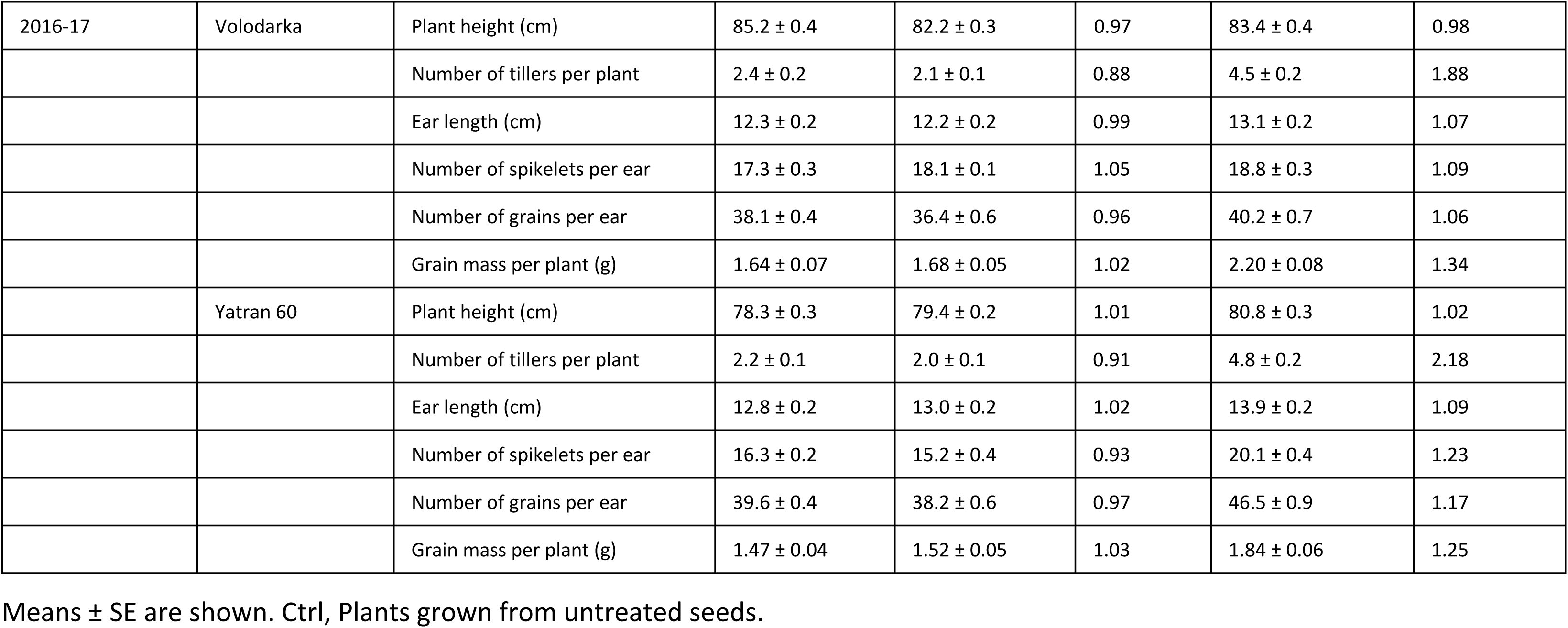
Productivity structure (2015-16 & 2016-17).

| Season | Wheat variety |  | Ctrl seeds | Pelleted seeds (P) | P/Ctrl ratio | Pelleted & Primed seeds (Pr) | Pr/Ctrl ratio |
| --- | --- | --- | --- | --- | --- | --- | --- |
| 2015-16 | Volodarka | Plant height (cm) | 93.1 ± 0.3 | 91.6 ± 0.6 | 0.98 | 94.3 ± 0.3 | 1.01 |
| | | Number of tillers per plant | 6.6 ± 0.1 | 7.1 ± 0.1 | 1.08 | 11.8 ± 0.2 | 1.79 |
| | | Ear length (cm) | 12.2 ± 0.2 | 11.9 ± 0.2 | 0.98 | 13.6 ± 0.1 | 1.12 |
| | | Number of spikelets per ear | 17.2 ± 0.2 | 18.4 ± 0.1 | 1.07 | 19.3 ± 0.1 | 1.12 |
| | | Number of grains per ear | 41.0 ± 0.5 | 42.5 ± 0.5 | 1.04 | 52.5 ± 0.6 | 1.28 |
| | | Grain mass per plant (g) | 1.57 ± 0.02 | 1.68 ± 0.02 | 1.07 | 2.18 ± 0.03 | 1.39 |
| | Yatran 60 | Plant height (cm) | 84.7 ± 0.4 | 83.3 ± 0.3 | 0.98 | 86.7 ± 0.3 | 1.02 |
| | | Number of tillers per plant | 7.3 ± 0.2 | 7.3 ± 0.1 | 1.00 | 12.7 ± 0.2 | 1.74 |
| | | Ear length (cm) | 13.4 ± 0.1 | 13.4 ± 0.1 | 1.00 | 13.5 ± 0.1 | 1.01 |
| | | Number of spikelets per ear | 21.0 ± 0.1 | 20.9 ± 0.1 | 1.00 | 21.6 ± 0.1 | 1.03 |
| | | Number of grains per ear | 44.0 ± 0.4 | 45.7 ± 0.3 | 1.04 | 55.8 ± 0.3 | 1.27 |
| | | Grain mass per plant (g) | 1.57 ± 0.03 | 1.68 ± 0.03 | 1.07 | 1.98 ± 0.02 | 1.26 |
| 2016-17 | Volodarka | Plant height (cm) | 85.2 ± 0.4 | 82.2 ± 0.3 | 0.97 | 83.4 ± 0.4 | 0.98 |
|  |  | Number of tillers per plant | 2.4 ± 0.2 | 2.1 ± 0.1 | 0.88 | 4.5 ± 0.2 | 1.88 |
|  |  | Ear length (cm) | 12.3 ± 0.2 | 12.2 ± 0.2 | 0.99 | 13.1 ± 0.2 | 1.07 |
|  |  | Number of spikelets per ear | 17.3 ± 0.3 | 18.1 ± 0.1 | 1.05 | 18.8 ± 0.3 | 1.09 |
|  |  | Number of grains per ear | 38.1 ± 0.4 | 36.4 ± 0.6 | 0.96 | 40.2 ± 0.7 | 1.06 |
|  |  | Grain mass per plant (g) | 1.64 ± 0.07 | 1.68 ± 0.05 | 1.02 | 2.20 ± 0.08 | 1.34 |
|  | Yatran 60 | Plant height (cm) | 78.3 ± 0.3 | 79.4 ± 0.2 | 1.01 | 80.8 ± 0.3 | 1.02 |
|  |  | Number of tillers per plant | 2.2 ± 0.1 | 2.0 ± 0.1 | 0.91 | 4.8 ± 0.2 | 2.18 |
|  |  | Ear length (cm) | 12.8 ± 0.2 | 13.0 ± 0.2 | 1.02 | 13.9 ± 0.2 | 1.09 |
|  |  | Number of spikelets per ear | 16.3 ± 0.2 | 15.2 ± 0.4 | 0.93 | 20.1 ± 0.4 | 1.23 |
|  |  | Number of grains per ear | 39.6 ± 0.4 | 38.2 ± 0.6 | 0.97 | 46.5 ± 0.9 | 1.17 |
|  |  | Grain mass per plant (g) | 1.47 ± 0.04 | 1.52 ± 0.05 | 1.03 | 1.84 ± 0.06 | 1.25 |

Means ± SE are shown. Ctrl, Plants grown from untreated seeds.

**Table 2.** Chlorophyll and carotenoid levels (2016-17).

| Wheat variety | Plant developmental stage | Chlorophyll a |  |  | Chlorophyll b |  |  | Carotenoids |  |  |
| --- | --- | --- | --- | --- | --- | --- | --- | --- | --- | --- |
|  |  | Ctrl seeds | Primed seeds (Pr) | Pr/Ctrl ratio | Ctrl seeds | Primed seeds (Pr) | Pr/Ctrl ratio | Ctrl seeds | Primed seeds (Pr) | Pr/Ctrl ratio |
| Volodarka | Flowering | 2.02 ± 0.01 | 2.27 ± 0.01 | 1.13 | 0.77 ± 0.01 | 0.77 ± 0.01 | 1.00 | 0.69 ± 0.01 | 0.69 ± 0.01 | 1.00 |
|  | Milk ripeness | 1.99 ± 0.01 | 2.13 ± 0.02 | 1.07 | 0.65 ± 0.02 | 0.72 ± 0.01 | 1.11 | 0.59 ± 0.01 | 0.65 ± 0.01 | 1.10 |
|  | Booting | 1.80 ± 0.01 | 2.11 ± 0.01 | 1.17 | 0.61 ± 0.01 | 0.77 ± 0.01 | 1.26 | 0.58 ± 0.01 | 0.61 ± 0.01 | 1.05 |
|  | Milk-wax ripeness | 0.97 ± 0.01 | 1.01 ± 0.01 | 1.04 | 0.30 ± 0.01 | 0.33 ± 0.01 | 1.10 | 0.52 ± 0.01 | 0.46 ± 0.01 | 0.89 |
| Yatran 60 | Flowering | 1.81 ± 0.01 | 2.61 ± 0.01 | 1.44 | 0.67 ± 0.01 | 0.94 ± 0.02 | 1.40 | 0.55 ± 0.01 | 0.77 ± 0.01 | 1.40 |
|  | Milk ripeness | 1.57 ± 0.01 | 2.44 ± 0.02 | 1.55 | 0.56 ± 0.02 | 0.83 ± 0.01 | 1.48 | 0.45 ± 0.01 | 0.68 ± 0.01 | 1.51 |
|  | Booting | 1.52 ± 0.09 | 1.91 ± 0.02 | 1.26 | 0.55 ± 0.03 | 0.62 ± 0.02 | 1.13 | 0.44 ± 0.02 | 0.55 ± 0.02 | 1.25 |
|  | Milk-wax ripeness | 1.10 ± 0.02 | 1.43 ± 0.01 | 1.30 | 0.31 ± 0.01 | 0.41 ± 0.01 | 1.32 | 0.45 ± 0.01 | 0.58 ± 0.01 | 1.29 |

Chlorophyll and carotenoids were measured as mg/g of fresh plant weight. Means ± SE are shown. Ctrl, Plants grown from untreated seeds.

**Fig 6.**
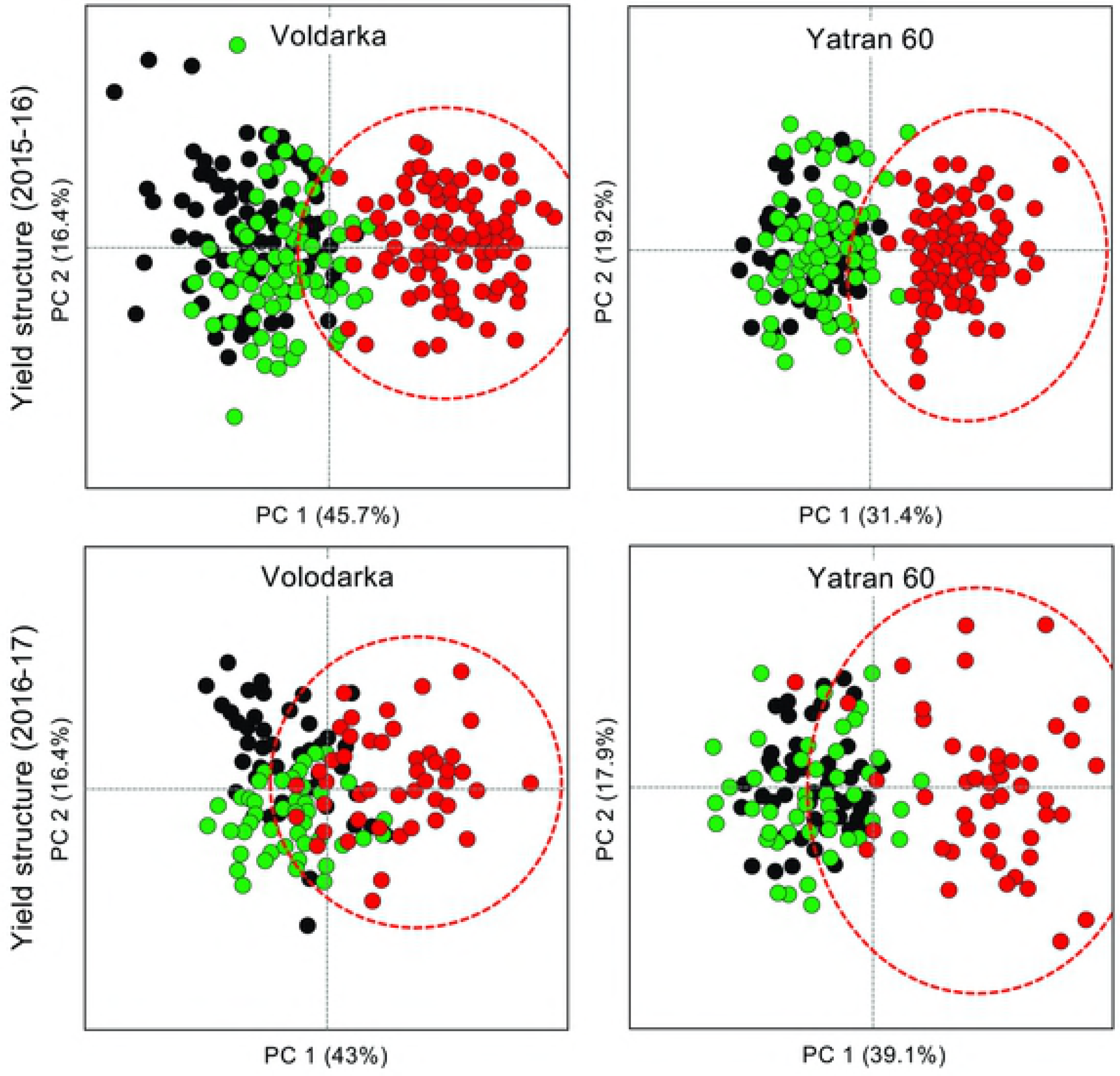
C6-HSL seed priming affects crop yield structure. The effect of seed treatment on winter wheat yield structure (over-all plant height, tiller numbers, ear length, spikelets per ear, grains per ear, and the grain mass per plant) can be visualised by Principal component analysis (PCA) for 2015 – 16 and 2016 – 17. Plants were grown from pelleted and primed (red and dashed oval), pelleted (green) or untreated (black) seeds. Axis scales are the same for each biplot and the cumulative percentage covariance is shown for both principal axes.

### Grain quality

Grain produced in 2016-17 was examined to determine whether seed priming effected grain quality or mean weight (**Fig 7**). Priming had a significant effect on grain quality with the proportion of good grain 1.3x greater than the untreated controls (p < 0.0001), though no wheat variety effects were observed (p = 0.08). A more complex range of factors including grain quality, seed treatment, and the seed quality × seed treatment and seed treatment × wheat variety interactions had significant effects on mean grain weight (p < 0.0001, 0.0164, 0.0002, 0.0006, respectively), though no wheat variety effect was observed (p = 0.46). These findings suggest that seed priming also effects grain quality and weight.

**Fig 7.**
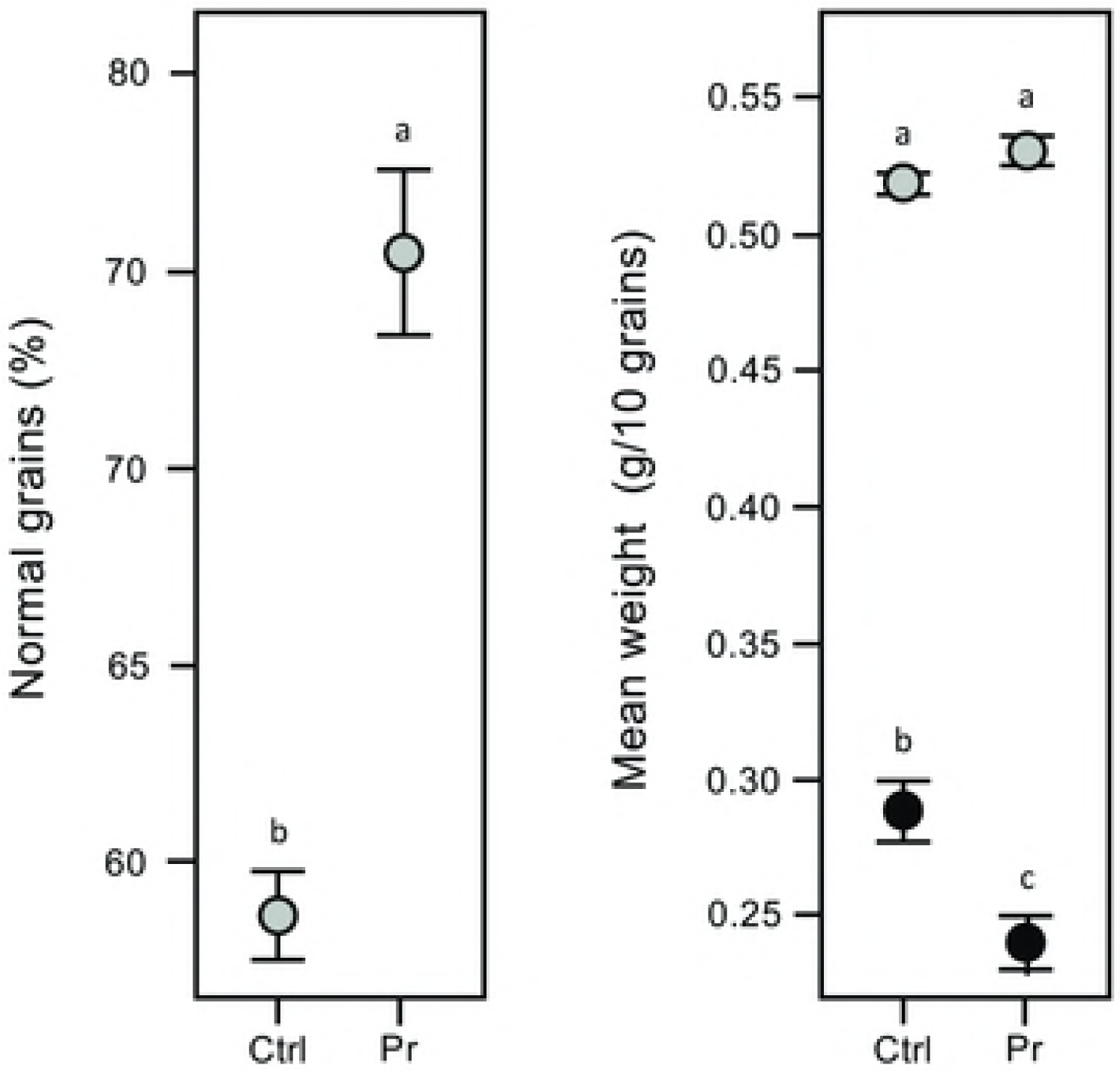
C6-HSL seed priming affects grain quality. Seed treatments have small but significant effects on the percentage of normal grain (grey) and on the mean weight of poor grain (black) in winter wheat crop trials (2016-17). Grain was collected from plants grown from pelleted and primed (Pr) or untreated seeds (Ctrl). As no significant wheat variety effect was observed (proportion of normal grain, p = 0.08; weight of 10 grains, p = 0.46) data are pooled and means and standard errors (SE) shown. Means not connected by the same letter are significantly different (α = 0.05).

### F1 plants grown from primed seed-plants

F1 grain collected from primed seed-plants grown in 2015-16 was tested to determine whether the seed priming effect was transferred to the next generation of wheat. Re-analysis of earlier experiments including F1 data showed significant differences between plants grown from F1 seeds and untreated control seeds in seven of 11 assays (**Table 3**) and this is also evident by PCA of crop yield data for 2016-17 by variety (**Fig 8**). This suggests that the seed priming effect is transferred to the following generation of wheat.

**Table 3.**
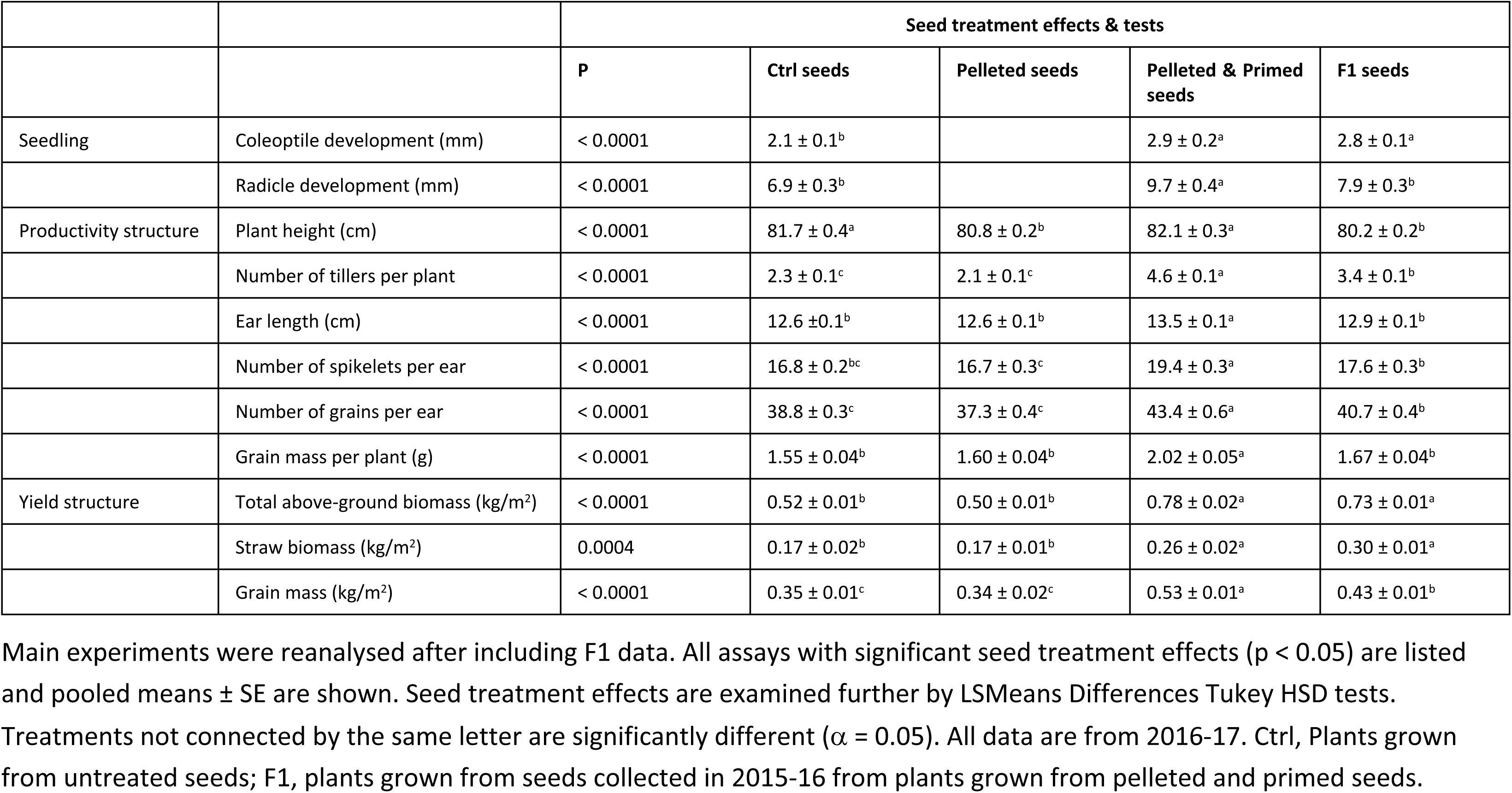
F1 characteristics (2016-17).

**Fig 8.**
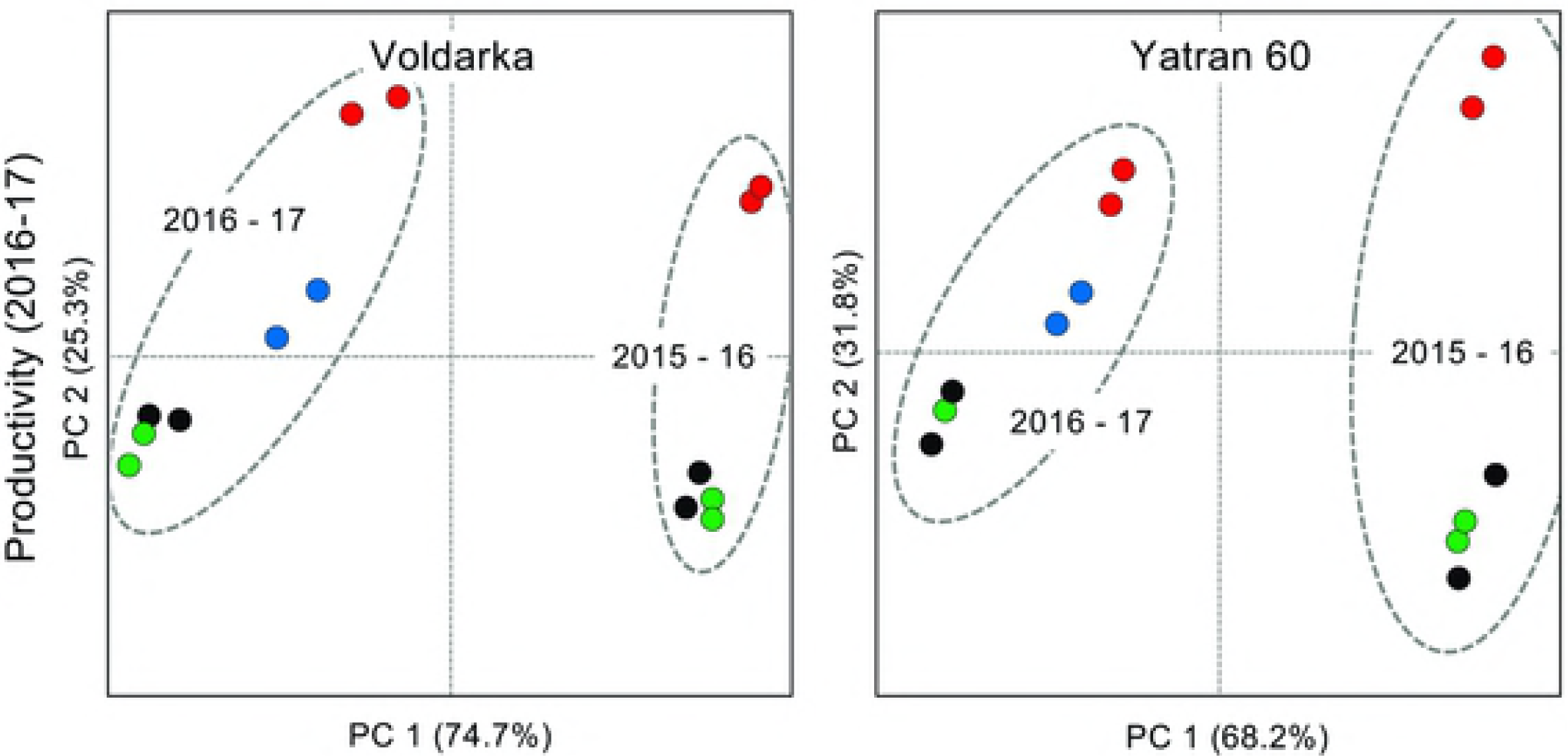
C6-HSL seed priming affects are transferred to the following generation. Principal component analysis (PCA) of winter wheat crop productivity (total above ground biomass, straw biomass, and grain mass) can differentiate plants grown from F1 seeds produced by pelleted and primed plants in 2015 – 16 and sown in 2016 – 17 (green) from plants grown from pelleted and primed (red), pelleted (green) or untreated (black) seeds. In these analyses a strong *year* effect is also apparent for both wheat varieties (indicated by dashed ovals). Axis scales are the same for each biplot and the cumulative percentage covariance is shown for both principal axes.

Main experiments were reanalysed after including F1 data. All assays with significant seed treatment effects (p < 0.05) are listed and pooled means ± SE are shown. Seed treatment effects are examined further by LSMeans Differences Tukey HSD tests. Treatments not connected by the same letter are significantly different (α = 0.05). All data are from 2016-17. Ctrl, Plants grown from untreated seeds; F1, plants grown from seeds collected in 2015-16 from plants grown from pelleted and primed seeds.

### Rhizosphere microbiology

Bacteria recovered from the roots of 2 month-old plants at the tillering stage in 2015-16 were characterised in a preliminary investigation to determine whether seed priming effected rhizosphere microbiology. Bacterial numbers in five functional groups (amylolytic, denitrifying, nitrifying, N-fixing, and oligonitrophilic bacteria) determined from pooled samples (**Table 4**) were found to be dependent on seed treatment for both wheat varieties (p < 0.0001). This preliminary result suggests that the seed priming effect alters the functional composition of bacterial communities around wheat roots.

**Table 4.** Rhizosphere microbiology (2015-16).

| Functional group | Volodarka |  |  | Yatran 60 |  |  |
| --- | --- | --- | --- | --- | --- | --- |
|  | Ctrl seeds | Pelleted seeds | Primed seeds | Ctrl seeds | Pelleted seeds | Primed seeds |
| Amylolytic | 11 | 32 | 120 | 10 | 33 | 23 |
| Denitrifying | 6 | 60 | 250 | 60 | 600 | 1 |
| Nitrifying | 9 | 190 | 19 | 500 | 1900 | 900 |
| N-fixing | 95 | 450 | 250 | 450 | 1500 | 150 |
| Oligonitrophilic | 230 | 480 | 270 | 440 | 930 | 1500 |

Numbers are $10^4$ per un-replicated (pooled) rhizosphere samples. Bacterial numbers and treatments are dependent for both wheat varieties (Volodarka, $\chi^2 = 568$ , $p < 0.0001$ ; Yatran 60, $\chi^2 = 1675$ , $p < 0.0001$ ). Ctrl, rhizosphere samples from plants grown from untreated seeds.

## Discussion

Early investigations of the effect of AHL applications on plant development and systemic resistance have focussed on *in vitro* and pot-plant experiments and reported changes in gene expression patterns, root development and systemic resistance [15-16, 28-32]. In this study, we were interested in determining whether bacterial AHLs can be used as a novel class of phyto-stimulator seed primers to improve cereal crop growth and grain yields, using winter wheat to investigate changes in seed germination *in vitro* and field trials to investigate changes in plant development and crop yield. Interestingly, the changes we have observed in C6-HSL-primed plants correspond well with those observed in wheat plants treated with the gram-negative plant growth-promoting rhizobacteria [33-35] which naturally produce AHLs to regulate gene expression and coordinate social activities [16, 36-38].

In this study, we investigate the effect of AHL applied directly to seeds before germination whereas previous research has focused on dosing or spaying seedlings or even older plants with AHL solutions [15-16, 30-32, 39-40]. We test here the effectiveness of seed priming with C6-HSL on two varieties of winter wheat as we expected to see a strong genetic effect in response to AHL treatment, but variety effects were seen in less than half of the characteristics we measured. The changes we observed in C6-HSL-primed plants suggest that the delivery of AHLs by seed priming would be successful in an agricultural context, as priming and pelleting are commonly used to apply a range of bactericides, fungicides, pesticides, herbicides, and phytostimulators, as well as fertiliser, to improve crop yields and lessen waste in a cost-effective manner.

### Effect of seed priming with C6-HSL on plant development and crop yield

We have investigated the effect of seed priming with C6-HSL on the germination and early seedling development of winter wheat *in vitro* where C6-HSL was found to have a small but significant positive effect on the germination and early seedling development (**Fig 4 & Table 3**). Seed priming with C6-HSL continued to have a positive effect in field trials, with improvements seen in plant biomass at the tillering stage (**Fig 5**), chlorophyll and carotenoid levels (**Table 2**), and on plant structure and yield at maturity (**Fig 7, Tables 1 & 3**). Collectively seed priming with C6-HSL seems to produce more robust and healthy plants which results in a higher grain yield at maturity. Although we have not directly investigated whether seed priming improves plant resistance to stress, we note that the 2016-17 growing season with lower than normal seasonal rainfall (**Fig 3**) resulted in plants with reduced productivity compared to 2015-16, yet grain yield from plants grown from primed seeds remained similar in both seasons (**Table 1**). The impact of seed priming on plant health and yield might be the result of direct effects on plant development and stress resistance and/or indirect effects resulting from changes in the plant-associated microbial community.

### Direct effects on plant development and stress resistance

Plants may respond directly to bacterial AHLs as they produce structurally-similar compounds including *N*-acyl ethanolamines and alkamides with likely roles in cell division and differentiation processes [28]. Bacterial AHLs are furthermore known to alter cell division in the primary root meristem, interact with G-protein and Ca^2+^ signalling, effect transpiration, and involve modulators of plant defence such as oxylipin and salicylic acid [8-9, 28, 30-31, 41], and in the context of pathogen and stress resistance, produce healthier and more robust plants. From our analyses, C6-HSL priming appears to promote faster germination and embryo development in winter wheat (**Fig 4**) which might allow seedlings to out-pace or overcome stresses which normally limit growth and yields. Alternatively, C6-HSL might effect signalling pathways to produce a response similar to the stress-induced morphogenic response (SIMR) caused by exposure to chronic, low-intensity stress [42-44]. It is possible that such ‘pre-stressed’ plants are better suited to subsequent growth under non-optimal conditions, and that this explains the positive impact on winter wheat plant development and yield we observed in our field trials.

We note that plants grown from seed collected from C6-HSL treated plants (i.e. second generation or F1 plants) performed significantly better than control plants (**Tables 8 & 9**). Although these improvements may partially be explained by healthy plants producing better quality seeds leading to more vigorous plants in the next generation [45-46], it is also possible that the seed priming effect is acting through epistasis in which heritable changes in gene expression patterns occurring without alteration of DNA sequences [47-48] and such epigenetic changes are known to occur in plants development and as a result of environmental stress [49].

### Indirect effects on plant-associated microbial communities

Rhizosphere (root) and phyllosphere (above ground) plant-associated microbial communities are known to have a direct and positive impact on plant growth and resistance to pathogens by mobilising soil nutrients, out-competing pathogens and inducing plant resistance [50-51]. It is possible that the C6-HSL carried by primed seeds might leach into the surrounding soil and stimulate plant growth-promoting (PGP) bacteria including *Bacillus* and *Pseudomonas* spp., resulting in better plant–surface colonisation and greater levels of pathogen suppression, especially during the early stages of seedling development. However, we believe that it is more likely that the direct stimulation of plant growth by C6-HSL results in greater levels of photosynthate exudate which then supports more effective PGP microbial communities. Our preliminary analysis of wheat rhizosphere community structure indicates that seed priming with C6-HSL results in functional changes (**Table 4**). We noted an increase in amylolytic bacteria which are involved in the degradation of dead root cells that promotes root growth and releases sugars for other rhizosphere bacteria, and they also effect plant defence response by provoking systemic resistance and improving the recognition of pathogenic microbes [52]. We also noted an apparent reduction in N-fixing and nitrifying bacteria which sometimes results in lower yields [53] though it might also be caused by improved N uptake by plants with better root development.

### Agro-economic advantages to AHL seed priming

Plant protection is ranked high in importance for stable wheat production by growers who face a range of pathogens which can be responsible for 15 – 20% yield losses per annum [54-55]. However, the net returns from fungicide use are highly variable and are only profitable in years with moderate to high disease severity [56-57]. As a consequence, the adoption of a new phyto-stimulator such as C6-HSL by growers will be governed by yield (and yield stability) improvements, chemical and treatment costs, and any potential savings made by reduced bactericide and fungicide applications.

We estimate our treatment costs at €14 per 10^3^ kg of seeds having synthesized C6-HSL using standard laboratory-grade reagents (analytical-grade C6-HSL costs €11 mg^-1^), and could probably lower costs even further using Ukrainian-sourced chemicals. Seed priming of winter wheat in Ukraine is increasing, with more that 70% of seeds treated with 1.5 × 10^9^ kg of fungicides in 2018 (Superagronom.com), but in order for C6-HSL seed priming to be commercially acceptable, costs need to be equal or lower than current treatments and produce the same or greater benefits. In comparison, the phyto-stimulator Raykat (Start Atlantica, Spain) and the fungicide Vitavax 200 FF (Brook Crompton, Italy), both widely used in Ukraine, cost €26 and €14.5 per 10^3^ kg of seeds, respectively. The continued use and development of bactericides, fungicides, and phyto-stimulators, etc., is necessary to increase stable crop yields world-wide, reduce the ecological impact of modern intensive agriculture, and combat the spread and evolution of new pathogens driven by globalisation and climate change [56, 58-59].

## Conclusions

Bacterial AHLs such as C6-HSL are known to effect plant growth and resistance to pathogens in laboratory-based experiments. Our work provides the first field-scale evidence to suggest that C6-HSL may be a viable phyto-stimulator when applied as a seed primer for winter wheat. The significant improvements seen in wheat growth, productivity and yield structure are explained by healthier and more robust plants resulting from the direct or indirect effects of C6-HSL on seed germination, plant growth and maturation. The development of new phyto-stimulators based on bacterial AHLs as a part of modern plant protection, in a combination with new bactericides, fungicides, crop varieties, and agricultural techniques, etc., could provide a more sophisticate means of combating plant pathogens and increasing stable crop yields in the future.

## Acknowledgments

We thank the Institute of Molecular Biology and Genetics, National Academy of Science of Ukraine, and Abertay University, for their continued support of the on-going collaboration between O.V. Moshynets and A.J. Spiers; A.J. Spiers is also member of the Scottish Alliance for Geoscience Environment and Society (SAGES). We thank the Director of the Institute of Plant Physiology and Genetics of the National Academy of Sciences of Ukraine, Prof. DSc. Academician V.V. Morhun for his support with the seed materials and fruitful discussions of our experimental results. We also thank the founder of Agrohub (https://agrohub.org/) and co-founder of Radar Tech (http://radartech.com.ua/) Yulia Poroshenko for her support and advice.

## Author Contributions

XXX

